# Museomic approaches to genotype historic *Cinchona* barks

**DOI:** 10.1101/2022.04.26.489609

**Authors:** Nataly Allasi Canales, Elliot M. Gardner, Tobias Gress, Kim Walker, Vanessa Bieker, Michael D. Martin, Mark Nesbitt, Alexandre Antonelli, Nina Rønsted, Christopher J. Barnes

## Abstract

Over the last few centuries, millions of plant specimens have been collected and stored within herbaria and biocultural collections. They therefore represent a considerable resource for a broad range of scientific uses. However, collections degrade over time, and it is therefore increasingly difficult to characterise their genetic signatures. Here, we genotyped highly degraded *Cinchona* barks and leaves from herbaria using two separate high-throughput sequencing methods (HtS) and compared their performance. We subsequently genotyped specimens using genome skimming, the most commonly performed high-throughput sequencing (HtS) technique. We additionally used a recently developed capture bait set (Angiosperm353) for a target enrichment approach. Specifically, phylogenomic analyses of modern leaf and historical barks of *Cinchona* were performed, including 23 historical barks and six fresh leaf specimens. We found that samples degraded over time, which directly reduced the quantity and quality of the data produced by both methodologies (in terms of reads mapped to the references). However, we found that both approaches generated enough data to infer phylogenetic relationships, even between highly degraded specimens that are over 230 years old. However, the target capture kit produced data for target nuclear loci and also chloroplast data, which allowed for phylogenies to be inferred from both genomes, whereas it was only possible to use chloroplast data using genome skimming. We therefore find the Angiosperms353 target capture kit a powerful alternative to genome skimming, which can be used to obtain more information from herbarium specimens, and ultimately additional cultural benefits.

## Introduction

Plant specimens have been stored in natural history collections for centuries and represent a major resource for understanding the life histories of extinct and extant species (Gutaker and Burbano, 2017). Such collections include type specimens, which are the original reference plant material that usually possess the defining features of that taxon used for the description of a species. These specimens have been used to build taxonomic systems, and to infer relationships and evolutionary histories among species. However, solely relying on the morphology of type specimens has led to inaccurate phylogenetic conclusions since underlying genomic variation may not lead to observable morphological traits (Goodwin et al., 2015), especially as some type specimens lack defining morphological features or are inaccurately annotated. Genome-level DNA data is being increasingly used in conjunction with morphological characters, and is becoming the standard in evolutionary studies. However, over time, the DNA molecules of specimens within herbaria become increasingly degraded; and it has been estimated that degradation may be six times faster for leaf material than for ancient bones (Weiß et al., 2016). This creates technical challenges that need to be resolved if historical plant specimens are to be utilised in both morphological and genomic analyses.

Specialised methods and analyses have been developed to process highly degraded DNA, allowing us to overcome some of the challenges of working with historical plant specimens within herbaria and elsewhere (Canales et al., (*in press*) ; Kistler et al., 2020). One of the first molecular studies of palaeobotanical remains used a CTAB DNA extraction of several tissue types and species, which allowed genetic information to be retrieved from specimens that were up to 44,600 years old (Rogers and Bendich, 1985). While PCR-based methods have been the norm (Drábková et al., 2002; Saltonstall, 2002; Jankowiak et al., 2005; Akhmetzyanov et al., 2020), they offered no means of authenticating the sequencing data from the sample (i.e. endogenous DNA rather than contamination). Now historical DNA (hDNA) studies have progressed to high-throughput sequencing (HtS), allowing for much larger portions of the genome to be sequenced from the many small DNA fragments (Palmer et al., 2012). The adaptation of HtS methods to sequence degraded DNA allows for further uses of herbarium vouchers, with a large proportion of herbarium material able to be utilised via these approaches. Sequencing methods are particularly relevant to specimens that do not include the morphological traits that allow for taxonomic identification, such as flowers, fruits and leaves. Sequencing methods therefore allow for phylogenetic inference and species annotation of bark and wood specimens that may otherwise have not been utilised (Bieker and Martin, 2018).

Previous studies on historical plant DNA that have used genome skimming approaches for systematics showed highly repetitive genomic regions like organelle genomes and rDNA can be recovered (Straub et al., 2012). They focused on analysing several types of historical specimens like pollen, and waterlogged pips (Gómez-Zeledón et al., 2017; Ramos-Madrigal et al., 2019). Few studies have been performed on ancient wood, which is generally considered to contain very low amounts of ultra-short and fragmented DNA (Dumolin-Lapègue et al., 1999). However, to our knowledge historical barks have not been explored until now. These pose an extra challenge as parts of the bark tissue are already dead on living trees (Bamber and Fukazawa, 1985). Additionally, bark and wood samples have higher contents of metabolites that inhibit DNA extraction and library preparation steps of both stored and fresh barks (Deguilloux et al., 2002). Consequently, methods targeting specific loci of the genome (such as PCR or microsatellites) have produced low quality data, which is not always reproducible (Tani et al., 2003; Liepelt et al., 2006; Tang et al., 2011). This may have contributed to the lack of studies using historical bark and wood specimens.

Genome skimming is an untargeted, low-coverage HtS technique, enabling many small fragments of high-copy DNA to be sequenced, mostly of organellar genomes and repetitive nuclear regions (Twyford and Ness, 2017). HtS approaches have helped overcome some of the challenges of working with highly degraded DNA from wood, and it has become the standard methodology to reconstruct the plastid genome or to infer haplotypes by calling SNPs (Bakker, 2017). HtS methods have been successfully employed to produce large quantities of sequencing data from wood material up to 9,800 years old (Tani et al., 2003; Liepelt et al., 2006; Wales et al., 2016; Wagner et al., 2018). Alternatively, target capture is a HtS method that amplifies pre-selected orthologous loci across the genome (Albert et al., 2007; Gnirke et al., 2009), avoiding contaminants from microbes and even other plants that may have accumulated on specimens over many years (Bieker et al., 2020). It has been successfully employed on highly degraded historical plant material, such as maize kernels (Avila-Arcos et al., 2011), maize cobs (da Fonseca et al., 2015), ragweed herbarium material (Sánchez Barreiro et al., 2017), waterlogged grape pips (Ramos-Madrigal et al., 2019), and potato herbarium specimens (Gutaker et al., 2019) but has not yet been applied to woody samples.

Overall, baits have been developed to target phylogenetically informative single-copy loci (target capture) that have been exploited to provide robust patterns of evolutionary relationships. However, it is a highly resource intensive method, with considerable time and resources required in designing and synthesising custom baits (Andermann et al., 2019). Recently, the Angiosperms353 kit was developed as a standardised set of baits that can be used to perform phylogenetic studies across the angiosperms without this initial resource input (Johnson et al., 2019) and its effectiveness in working with herbarium material is being explored (Brewer et al., 2019). However, it has not been tested on highly degraded museum material yet.

In this study we focus on the *Cinchona* genus in the coffee family (Rubiaceae), which is of great cultural importance to the people of the Andean countries (Prendergast and Dolley, 2001). Due to the high alkaloid content in their barks, three *Cinchona* species have been of particular interest for wild harvesting and cultivation: *C. calisaya* Wedd. (yellow bark), *C. officinalis* L. (pale bark) and *C. pubescens* Vahl (red bark), as well as hybrids with *C. officinalis* (Nair and Prabhakaran Nair, 2010).

*Quinine-type alkaloids from the barks of the Cinchona* genus served as the main treatment for malaria for three hundred years from circa 1630s onwards. These compounds were first isolated as the active ingredient in 1820 (Thompson, 1928). During this period, interest increased in the potential cultivation of *Cinchona* to control the mass-production of quinine for European imperial projects, and plantations were established from the 1850s in tropical countries such as India, Indonesia, and Jamaica. Quinine remained the main treatment for malaria until its gradual replacement by synthetic antimalarials in the 1940s (Walker and Nesbitt, 2019). Quinine and other quinoline alkaloids are primarily located in the bark (around 3-12% of total biomass; McCalley, 2002, where they likely also have inhibitory activity in DNA extraction and library preparation steps (Deguilloux et al., 2002).

Today, there is a large historical resource of over 1,000 commercial barks samples, housed in the Economic Botany Collection of the Royal Botanic Gardens, Kew (RBGK), UK. A part of these samples have been identified to species level and have provenance data indicating they were collected across the late 18th and the late 19th centuries (Table 1; Canales et al., 2020). A better understanding of their history could enable the mapping of historical trade routes and better understanding of the chemical ecology of quinine alkaloids. Because of the difficulty of designating species of specimens such as these using a conventional morphology approach, palaeogenomics can be exploited to do this to ultimately maximise their cultural value. However, the DNA within these bark specimens have accumulated *post-mortem* damage through ageing, while retaining their alkaloids that potentially inhibit downstream amplification (Canales et al., 2020). Therefore generating informative genetic data from historical *Cinchona* barks is challenging.

**Table 1.** Target capture (TC) and high-throughput (HtS) results for modern and historical specimens of the *Cinchona* genus.

|  | Samples | Collection date | Country | Species | Name on record | Type of data | Raw reads | Reads mapped | Estimated coverage (X) | mapped to ref (%) | Age |
| --- | --- | --- | --- | --- | --- | --- | --- | --- | --- | --- | --- |
| 0 | Ccalisaya71 | 1876 | Indonesia | *C. calisaya* | *Cinchona ledgeriana* | HtS | 16,994,830 | 315,051 | 55.0 | 1.85 | Historical |
| 1 |  |  |  |  |  | TC | 29,000,734 | 2,202,016 | NaN | 7.60 | Historical |
| 2 | Ccalisaya34 | Post 1850 | Indonesia | *C. calisaya* | *Cinchona ledgeriana* | HtS | 9,587,571 | 167,214 | 41.9 | 1.74 | Historical |
| 3 |  |  |  |  |  | TC | 13,107,034 | 1,440,236 | NaN | 11.00 | Historical |
| 4 | Ccalisaya48 | 1867 | India | *C. calisaya* | *Cinchona pahudiana* | HtS | 13,588,982 | 649,320 | 86.1 | 4.78 | Historical |
| 5 |  |  |  |  |  | TC | 37,032,164 | 2,820,716 | NaN | 7.60 | Historical |
| 6 | Clancifolia28 | unknown 19th C | South America | *C. lancifolia* | *Cinchona lancifolia* | HtS | 11,920,430 | 110,019 | 24.3 | 0.92 | Historical |
| 7 |  |  |  |  |  | TC | 8,579,138 | 673,885 | NaN | 7.90 | Historical |
| 8 | Ccalisaya9 | unknown 19th C | South America | *C. calisaya* | *Cinchona calisaya* | HtS | 12,648,238 | 178,520 | 24.8 | 1.41 | Historical |
| 9 |  |  |  |  |  | TC | 34,401,668 | 2,880,128 | NaN | 8.40 | Historical |
| 10 | Cofficinalis101 | 1777-1788 | Peru | *C. officinalis* | *Cinchona officinalis* L. | HtS | 2,800,160 | 70,700 | 21.2 | 2.52 | Historical |
| 11 |  |  |  |  |  | TC | 3,480,514 | 293,044 | NaN | 8.40 | Historical |
| 12 | Cpubescens98 | 1777-1788 | Peru | *C. pubescens* | *Cinchona pubescens* var. *purpurea* or *ovata* | HtS | 7,734,184 | 49,305 | 16.9 | 0.99 | Historical |
| 13 |  |  |  |  |  | TC | 14,656,153 | 841,409 | NaN | 5.80 | Historical |
| 14 | Ccalisaya12 | 1873 | South America | *C. calisaya* | *Cinchona calisaya* Weddell | HtS | 8,123,059 | 39,494 | 5.3 | 0.49 | Historical |
| 15 |  |  |  |  |  | TC | 3,776,222 | 297,463 | NaN | 7.90 | Historical |
| 16 | Cglandulifera27 | 1854 | South America | *C. glandulifera* | *Cinchona glandulifera* | HtS | 12,791,708 | 376,195 | 65.7 | 2.94 | Historical |
| 17 |  |  |  |  |  | TC | 39,652,882 | 3,264,504 | NaN | 8.20 | Historical |
| 18 | Cmicrantha25 | 1873 | South America | *C. micrantha* | *Cinchona micrantha* | HtS | 7,392,343 | 545,697 | 86.4 | 7.38 | Historical |
| 19 |  |  |  |  |  | TC | 37,417,695 | 2,943,274 | NaN | 7.90 | Historical |
| 20 | Ccalisaya67 | 1838 | South America | Spurious cinchona | Spurious cinchona | HtS | 77,163,465 | 2,051,211 | 125.7 | 2.66 | Historical |
| 21 |  |  |  |  |  | TC | 86,057,272 | 6,524,964 | NaN | 7.60 | Historical |
| 22 | Spruceseed1861 | 1861 | Peru | *C. pubescens* | *Cinchona pubescens* | HtS | 13,298,661 | 169,066 | 84.0 | 6.05 | Historical |
| 23 |  |  |  |  |  | TC | 45,456,459 | 5,311,261 | NaN | 11.70 | Historical |
| 24 | Cpubescens84 | 1883 | India | *C. pubescens* | *Cinchona pubescens* | HtS | 8,413,518 | 85,520 | 32.8 | 1.61 | Historical |
| 25 |  |  |  |  |  | TC | 27,608,912 | 2,285,958 | NaN | 8.30 | Historical |
| 26 | Cnitida40 | 1852 | South America | *C. nitida* | *Cinchona nitida* | HtS | 26,708,967 | 178,482 | 32.5 | 0.67 | Historical |
| 27 |  |  |  |  |  | TC | 42,790,993 | 2,047,385 | NaN | 4.80 | Historical |
| 28 | Clancifolia33 | 1853 | Colombia | *C. lancifolia* | *Cinchona lancifolia* Mutis | HtS | 2,161,061 | 47,784 | 17.5 | 2.21 | Historical |
| 29 |  |  |  |  |  | TC | 14,442,875 | 1,624,012 | NaN | 11.20 | Historical |
| 30 | Ccalisaya3 | 1852 | South America | *C. calisaya* | *Cinchona amygdalifolia* Weddell | HtS | 16,850,139 | 186,755 | 27.1 | 1.11 | Historical |
| 31 |  |  |  |  |  | TC | 41,506,867 | 2,410,468 | NaN | 5.80 | Historical |
| 32 | Ccalisaya14 | 1853 | South America possibly Bolivia | *C. calisaya* Wedd. | *Cinchona calisaya* var *boliviana* | HtS | 6,171,873 | 22,144 | 4.4 | 0.36 | Historical |
| 33 |  |  |  |  |  | TC | 1,378,065 | 78,557 | NaN | 5.70 | Historical |
| 34 | Cofficinalis38 | 1852 | South America | *C. officinalis* L. | *Cinchona officinalis* L. | HtS | 9,093,075 | 163,218 | 37.1 | 1.79 | Historical |
| 35 |  |  |  |  |  | TC | 19,046,144 | 1,608,859 | NaN | 8.40 | Historical |
| 36 | Cangustifolia4 | 1854 | South America | *C. lancifolia* | *Cinchona angustifolia* | HtS | 47,277,018 | 876,880 | 82.9 | 1.85 | Historical |
| 37 |  |  |  |  |  | TC | 80,114,288 | 6,273,561 | NaN | 7.80 | Historical |
| 38 | Ciliosemina_pedunculata87 | unknown 19th C | South America | *Ciliosemina pedunculata* | *Remijia pedunculata* | HtS | 20,097,917 | 89,059 | 32.4 | 0.85 | Historical |
| 39 |  |  |  |  |  | TC | 46,751,177 | 4,532,440 | NaN | 9.70 | Historical |
| 40 | Cscrobiculata50 | 2014 | San Ignacio, Cajamarca, Perú | *C. scrobiculata* | *Cinchona scrobiculata* | HtS | 68,856,758 | 305,848 | 215.6 | 5.17 | Modern |
| 41 |  |  |  |  |  | TC | 2,183,034 | 228,280 | NaN | 10.50 | Modern |
| 42 | Cnitida51 | 2013 | San Ramón, Junín, Perú | *C. nitida* | *Cinchona nitida* | HtS | 8,552,790 | 10,324 | 5.6 | 5.09 | Modern |
| 43 |  |  |  |  |  | TC | 19,776,304 | 1,883,501 | NaN | 9.50 | Modern |
| 44 | Ccalisaya52 | 2014 | San Juan del Oro, Puno, Peru | *C. calisaya* | *Cinchona calisaya* | HtS | 37,180,062 | 154,391 | 84.5 | 5.22 | Modern |
| 45 |  |  |  |  |  | TC | 19,096,988 | 1,889,884 | NaN | 9.90 | Modern |
| 46 | Ccalisaya53 | 2014 | San Juan del Oro, Puno, Peru | *C. calisaya* | *Cinchona calisaya* | HtS | 37,611,592 | 171,736 | 112.5 | 5.65 | Modern |
| 47 |  |  |  |  |  | TC | 4,626,036 | 471,402 | NaN | 10.20 | Modern |
| 48 | Cpubescens54 | 2010 | Cusco, Perú | *C. pubescens* | *Cinchona pubescens* | HtS | 50,141,352 | 185,025 | 74.2 | 5.05 | Modern |
| 49 |  |  |  |  |  | TC | 4,418,265 | 478,796 | NaN | 10.80 | Modern |
| 50 | Cpitayensis55 | 2010 | Ayabaca, Piura, Perú | *C. pitayensis* | *Cinchona pitayensis* | HtS | 54,083,970 | 217,428 | 116.7 | 5.08 | Modern |
| 51 |  |  |  |  |  | TC | 15,362,944 | 1,394,553 | NaN | 9.10 | Modern |

The present study has three major objectives. The first objective is to demonstrate that DNA can be sequenced from the highly degraded *Cinchona* barks to provide meaningful genomic characterisations. In order to do this, we developed a customised DNA extraction protocol for historical barks, which was followed by either genome skimming or target capture using a standardised bait kit (Angiosperms353 Kit). Our second objective is to compare the performance of the data produced by the two different approaches in terms of reads generated, reads mapped, cost, endogenous DNA content, and the estimated coverage for historical bark samples and silica-dried modern leaf. Our last objective is to estimate the evolutionary relationship between samples using each method, and then compare them in terms of phylogenetic robustness and placement of taxa.

## Material and methods

### Sample selection

We sampled 23 historical *Cinchona* bark specimens housed at the Economic Botany Collection, Royal Botanic Gardens, Kew. These samples were collected between the 1780s and 1876 from their native Andean forest ranges as well as from cultivations in British-Indian and Dutch-Indonesian colonial plantations. The samples are from seven species and have detailed temporal and spatial data described in historical labels, publications and letters linked to each specimen (Figure 1, Table 1). We collected ∼100 mg of each historical bark. To minimise the risk of contamination with non-degraded or amplified DNA, we performed all pre-PCR work in the dedicated, positively pressurised palaeo-genomic facilities at the GLOBE Institute, University of Copenhagen (Denmark). For comparison, we included six modern leaf samples representing five species, which were collected in 2014 in their native range within South America and were dried and stored in silica gel (Table S1). The voucher specimens are deposited in the Natural History Museum of Denmark, University of Copenhagen Herbarium (C).

**Fig 1.**
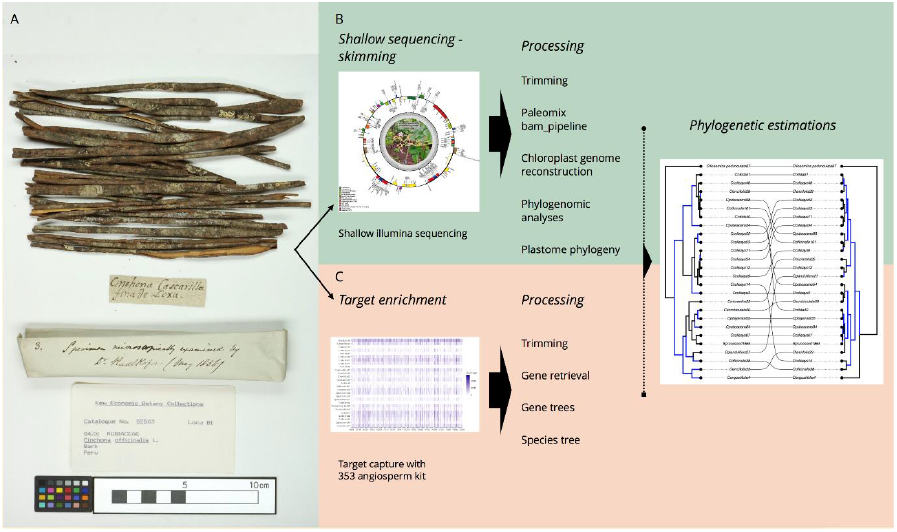
Historical bark sample and overview of the strategy (A) Study sample *C. officinalis* (Cofficinalis101, *C. condaminea* in the original label) from the Economic Botany Collection, Kew. (B) Chloroplast genome reconstruction using a reference genome. (C) Gene retrieval efficiency of the Angiosperms353 nuclear kit in historical barks and modern leaf samples.

### DNA extraction and initial library preparation

The historical samples underwent DNA extraction using a modified version of Wagner’s protocol for DNA extraction from wood (Wagner et al., 2018), customised to maximise endogenous DNA recovery using protocols developed for working with degraded DNA (Wales and Kistler, 2019). Briefly, our method includes these steps: (i) cleaning of the bark surface with sterile and previously UV-irradiated scalpels; (ii) grinding of ∼100 mg material in sterile mortars; (iii) overnight digestion of the bark at 37 °C with a lysis buffer made of 10 mM Tris-HCl, 10 mM NaCl, 2% w/v SDS, 5 mM CaCl2, 2.5 mM EDTA, 40 mM DTT, 2% PVP and 10% proteinase K; then we followed Wagner’s protocol precisely. The quality and quantity of DNA extracts were measured using a TapeStation 4200 instrument (Agilent Technologies, Germany) with High-Sensitivity ScreenTape D1000 reagents (Agilent Technologies, Germany). For the modern silica gel-dried samples, 20 mg of dried leaves underwent DNA extraction using a DNeasy plant mini kit (Qiagen, Valencia, CA, USA) as per manufacturer’s instructions.

All DNA extracts underwent initial library preparation until indexing, after which indexed libraries were divided into two and processed in separate ways. First, for both sample sets, BEMC blunt-end-multi column DNA ligation (Carøe et al., 2018) was performed using 32 μL of DNA input. We assessed the DNA library concentration through real-time PCR (Roche LightCycler 480) and SYBR Green chemistry (ThermoFisher Scientific, Denmark). Real-time PCR was conducted in 20 μL reaction using 1 μL of 1:20-diluted library, 0.1 U μL^-1^ TaqGold (Invitrogen, Life Technologies), 1x Buffer, 2.5 mM MgCl2, 1 mg mL^-1^ BSA, 0.2 mM dNTPs, 0.2 μM Illumina in PE1.0 primer (5′-AAT GAT ACG GCG ACC ACC GAG ATC TAC AC (index) T CTT TCC CTA CAC GAC GCT CTT CCG ATC T) and 0.2 mM Multiplex Index Primer (5′-CAA GCA GAA GAC GGC ATA CGA GAT (index) GTG ACT GGA GTT CAG ACG TGT GCT CTT CCG), where (index) are unique, 6-mer index tags. The amplification conditions were as follows: 10 min at 92 °C (initial activation by heating), 40 cycles of 30 s at 92 °C (denaturation), 30 s at 60 °C (annealing), and 30 s at 72 °C (elongation), and then 7 min at 72 °C (final elongation). Ct values were determined using the second-derivative maximum algorithm implemented in LightCycler software (Roche Applied Science, Germany). Index PCR was performed with the same conditions as real-time PCR, except that we (i) amplified two replicates per library which were pooled during the purification process, (ii) used 5 μL of the undiluted library, (iii) the number of amplification cycles was calculated by the C_T_ value minus 7, to correct for dilution effects. PCR products were purified using MagBio HighPrep PCR (MagBio Genomics, USA) magnetic beads and adding 1.8X (bead:DNA volume), in a final volume of 50 μL and were diluted to 1.5 ng uL^-1^ to be measured in the High-Sensitivity ScreenTape D1000 reagents (Agilent Technologies, Germany). We included blank controls in all laboratory processes together with our samples. They include extraction, library, PCR and indexing blank controls. Additionally, we indexed the blank controls with the maximum number of cycles from our samples even when there were non-detectable DNA quantities on a TapeStation instrument (Agilent Technologies, Germany).

### Two library preparation approaches: genome skimming and target capture

At this point, libraries were divided into halves that underwent two separate methods: 1) a standardised approach for shotgun sequencing to produce genome skimming data (Wagner et al., 2018) and 2) target capture using a standardised Angiosperms353 kit (Arbor Biosciences myBaits Target Sequence Capture Kit, ‘Angiosperms353 v1’ MI, USA; (Johnson et al., 2019)). For the genome skimming method, we pooled the libraries at equimolar concentration and sequenced them in a single lane on a HiSeq4000 instrument in SR80 mode at the Danish National High-Throughput Sequencing Centre (see Table S1 for more details) for historical samples. For the target capture method, after double-indexing the samples, the myBaits v.4.01 protocol was followed using the Angiosperms353 kit. For this, 8 reactions were used, with each reaction consisting of 12 µL of four indexed libraries. The incubating reaction of libraries and baits was equal to 30 μL and the hybridisation temperature was 65 °C for 18 h. The cleaning step was done with magnetic beads three times. While attached to beads, we amplified the libraries using the KAPA HiFi HotStart ReadyMix (Kapa Biosystems, Wilmington, MA, USA) with 14 PCR cycles to reach sufficient quantities for sequencing. After libraries were enriched, we pooled them in equimolar concentrations for sequencing on a HiSeq4000 lane (PE 150 mode) at the GeoGenetics Sequencing Core (University of Copenhagen, Denmark).

### Bioinformatic analyses of the genome skimming libraries

Genome skimming data were processed using Paleomix v1.2.5 (Schubert et al., 2014). Initially, the raw reads were trimmed and cleaned using AdapterRemoval2 (Schubert et al., 2016) and reads <30 bp were discarded. The collapsed resulting reads were then mapped against the reference chloroplast genome of *Cinchona pubescens* (157 Kb: Canales et al, unpublished data) using the Paleomix bam pipeline with the BWA-backtrack algorithm within the Burrows-Wheeler Aligner (Li and Durbin, 2009). Seeding was disabled as per recommendations from Schubert (Schubert et al., 2012), alignments with all mapping quality were taken into account, unmapped reads and PCR duplicates were filtered. Additionally, all historical data was authenticated by measuring the deamination pattern (C-to-T substitutions) using mapDamage2.0 (Jónsson et al., 2013). We obtained consensus plastid genome haplotypes from the bam files using Geneious Prime 2022.0.1 (https://www.geneious.com) (Anon, 2019), which were then aligned using MAFFT v 7.453 in the auto mode (Katoh et al., 2005). Finally, the phylogenetic inferences were performed using maximum-likelihood estimations within RAxML-NG v 1.0 (Kozlov et al., 2019) with the GTR+G model of molecular evolution, and the robustness of the consensus tree was tested by resampling 1000 bootstrap replicates.

### Bioinformatic analyses of the enriched libraries

While Paleomix has been successfully used with bait data (Sánchez Barreiro et al., 2017), our attempt to retrieve the enriched sequences with the same strategy as used for genome skimming yielded extremely low coverage. Conversely, using HybPiper, a tool for Hyb-Seq data (Johnson et al., 2016), a substantially higher number of reads were retrieved. First, raw reads were cleaned and trimmed using AdapterRemoval2 (Schubert et al., 2016), which were then processed using the HybPiper pipeline with default parameters, except for coverage which was set to be at least 2x, with the clean reads mapped to the target file (<u>Angiosperms353 - targetSequences</u>) using BLAST+ v. 2.9 (Madden, 2013). Contigs were assembled using SPAdes v. 3.13.1 (Bankevich et al., 2012) and aligned to the target sequence using Exonerate v. 2.4.0 (Slater and Birney, 2005). A heatmap was constructed using the ggheattree command in the treeheatr package in *R* (3.6.1) to visualise the gene recovery within samples. (Madden, 2013). Each gene set was aligned using MAFFT v.7.453 in the auto mode (Katoh et al., 2005). Then, the columns with > 75% gaps were removed. For historical and/or low-recovery specimens, poorly aligned ends were trimmed using HerbChomper Beta version 0.3 (Gardner, 2020). Long-branches were detected with TreeShrink v.1.3.9 (Mai and Mirarab, 2018) and the outlier sequences were removed using trimAl v1.2 (Capella-Gutiérrez et al., 2009). Phylogenetic trees were estimated using IQ-tree v2.1.2 (Minh et al., 2020) and the resulting trees’ branches corresponding to partitions that were reproduced in less than 30% of bootstrap replicates were collapsed using TreeCollapse4 v3.2 (Hodcroft, 2011). Then, all trees were concatenated in a supermatrix using HybPiper and the final phylogenetic tree was estimated using IQ-tree with model finder followed by tree inference and ultrafast bootstrap of 1000.

Finally, since the target capture approach focuses on retrieving nuclear genome regions, and the genome skimming mainly retrieves high-copy regions like the chloroplast genome, they can show different evolutionary histories. Therefore, we retrieved reads assigned to the chloroplast from target capture data and compared plastid phylogenetic trees for both methods. To retrieve plastid reads from the target capture data, reads were mapped against the plastid genome of *C. pubescens* using Paleomix’ bam_pipeline. Then, a consensus for plastid genomes was called N if the coverage < 5 using Geneious Prime 2022.0.1. Finally, the phylogenetic trees were estimated by aligning the consensus plastid genomes using MAFFT v7.453 in auto mode and inferring the tree with RAxML-NG v 1.0, bootstrap value of 1000 and GTR+G as substitution model.

### Statistical analyses

All statistical analyses were performed in the *R* (3.6.1) statistical computing environment. Differences in methodologies were compared using paired Student’s *t*-tests, with differences in reads produced, reads mapped and percent reads mapped. The effects of degradation (Appendix Fig. S1) were analysed by performing unpaired Student’s *t*-tests on the genome-skimming data and target-enrichment datasets separately. Samples were considered as either historical or modern, and the reads produced, reads mapped and reads mapped (%) were compared using unpaired Student’s *t*-tests. Meanwhile, plastid and nuclear trees were visualised and compared plotted in a tanglegram (Racine, 2012) using phytools (Revell, 2012). The Roubinson-Foulds metric was calculated to quantify the distance between phylogenetic trees, which was performed using the treediff function within the phangorn package in *R* (Schliep, 2011). Differences between trees were visualised as tanglegrams using the cophylo function, also within the phytools package (Revell, 2012).

## Results

### Generating endogenous DNA from historical barks

From the 30 historical specimens sampled, 26 had sufficient quality DNA to be sequenced, while all six of the modern silica-gel dried leaves underwent successful sequencing. We considered a sample for sequencing when after indexing it showed at least 100 pg μl^-1^ of DNA that was between 140 - 700 bp in length using a HighSensitivity analysis (TapeStation, Agilent Technologies, Germany). From these, we recovered genome skimming and target captured data from DNA extracts of 23 historical barks and six leaf samples spanning the *Cinchona* genus in South America and plantations in Asia (Appendix Tab. S1).

Regarding the genome skimming data, the mean number of reads produced per sample was 22,586,297 (+/-20,540,754), of which 285,246 (+/-405,276) reads mapped the plastid genome (Fig. 2). The average read length recovered after trimming was 60.8 bp for historical bark samples and 111.5 bp for modern leaf samples. The estimated coverage from unique hits for the historical samples ranged from between 4.3 and 125.7x, but for the modern leaves, coverage was significantly higher (*t* = -1.94, *P* = 0.103), ranging from 5.6 and 215.6x coverage. The percentage of reads from modern silica dried leaves that mapped to the plastid genome ranged from 5.1% - 5.7%, with Ccalisaya53 specimen yielding the highest percentage. For historical barks, the percentage of reads mapping the plastid genome is 0.35 - 7.38%. The highest endogenous content was for a historical bark collected in 1873 from South America (Cmicrantha25). The lowest was Ccalisaya14, collected in 1853 from South America, possibly from Bolivia (based on historical metadata labels). Additionally, there was a significant reduction in the number of reads in the historical specimens compared to the modern (*t* = -2.85, *P* = 0.024) and the percentage of reads mapped to the reference (*t* = -7.07, *P* < 0.001) but not in the number of mapped reads (*t* = 1.30, *P* = 0.208).

**Fig 2.**
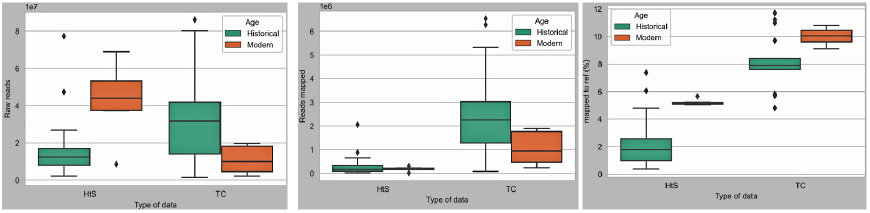
Overview of the samples per age, (A) raw reads generated, (B) reads mapped, (C) percentage of reads mapped to a reference, plastid genome and the 353 loci for the genome skimming (HtS) and target capture (TC) approaches, respectively.

For the enriched data, the mean number of reads produced per specimen was 26,908,832 (+/-22,152,659), of which 509,242 (+/-421,822) reads mapped to the updated 353 loci target file (McLay et al., 2021). The highest number of raw and mapped reads was generated for Ccalisaya67 collected in 1838 in South America. The percentage of reads mapped to the target for historical barks was between 4.8% and 11.7% (being Spruceseed1861 the highest), while for modern leaf samples it was between 9.10% and 10.8%.

`We recovered on average a total of 108 genes at ≥ 50% of the total length for historical samples, while for the modern samples, an average of 160 genes were recovered.` The percentage of reads that mapped in historical samples on average was 8.1%, while for the modern samples it was 10% (Table 1). Surprisingly, the specimen which had the longest gene length recovered was Ccalisaya67, a historical sample. From the paralog analyses, six samples had paralog warnings, with between one and six warnings in three historical samples, and between three and six warnings in three modern samples. We also attempted to retrieve the angiosperms genes from the genome skimming data, but the gene recovery was too low to be informative (below 0.1x coverage). Additionally, there was a significant effect of sample degradation, with fewer raw reads (*t* = 3.33, *P* = 0.003) and mapped reads (*t* = 2.79, *P* = 0.011) produced in the historical specimens compared to the modern, which corresponded to less endogenous DNA being recovered (*t* = - 4.00, *P* = 0.011).

The retrieval of authentic historical DNA (i.e. not modern contaminants) were authenticated with means of quantifying C-to-T substitutions and observed an excess of those at the first base on 5’ end of sequenced molecules (Appendix Fig. S1). Finally, the gene recovery efficiency ranged from 0.2% to 61.6% for historical samples, and between 18.4 to 54.2% for modern samples (Fig. 3). The number of target genes retrieved (sequences covering at least 50% of the target length) ranged from 1 to 229 for the historical barks. While for the silica gel-dried specimens were from 39 to 190.

**Fig. 3.**
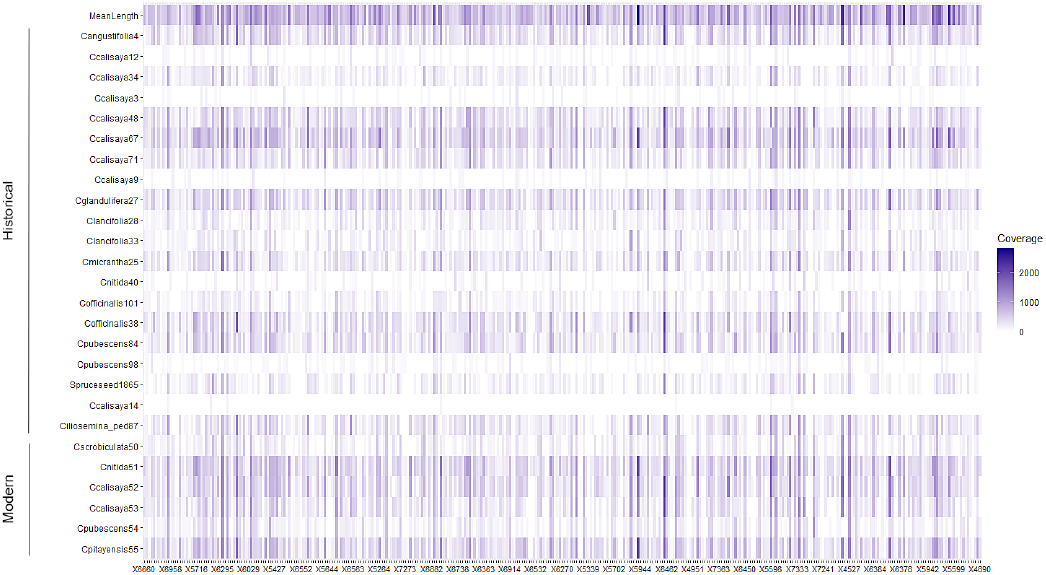
Heat-map showing the recovery efficiency of the Angiosperms353 kit loci for the historical bark samples and the silica gel-dried leaves of the *Cinchona* genus.

### Comparing the data production of the genome skimming and target capture

There were no significant differences in the number of raw reads produced with each method (*t* = -7.99, *P* = 0.432), with target capture producing a mean of 26,908,832 (+/-22,152,659) reads and the genome skimming producing a mean of 22,586,297 (+/-20,540,754) reads. However, there were significant differences observed between the mapped reads (*t* = -2.59, *P* = 0.016) and the percentage of reads mapped to target (*t* = -6.23, *P* < 0.001), with target capture producing significantly more of each (*t* = -15.23, *P* < 0.001). Within the target capture data, 509,242 (+/-421,822) mapped reads were produced, whereas 285,246 (+/-405,276) mapped reads were produced in the genome skimming approach. Similarly, the percentage of mapped reads using the genome skimming approach was 2.9 (+/-4.29) %. While for the target capture data was 8.53 (+/-3.24) (%).

Cost is another variable to consider when comparing methods. Using the target capture approach requires extra costs associated with the Angiosperms353 baits (approximately $1000 for 8 reactions in 2021) and KAPA HiFi HotStart ReadyMix (approximately $160 for 100 reactions in 2021), leading to a $40 extra for each sample for the target enrichment approach. Although the enrichment method with the Angiosperms353 kit produces more raw reads and mapped reads, it does not necessarily increase the recovery of ultra-short endogenous DNA.

### Comparison of the phylogenetic performance of HtS and target capture in Cinchona specimens

Phylogenetic trees were created using the differing methodologies. The HtS genome skimming data based on the plastid genome can be phylogenetically analysed as one unit with one unique evolutionary history (Appendix Fig. S1). Meanwhile, to analyse the multilocus nuclear data from target capture, a concatenation approach was used prior to inferring a nuclear tree (Appendix Fig. S3). In order to compare the two trees, we visualised them in a tanglegram revealing different topologies (Fig. 4A). The Robinson-Foulds distance between trees was calculated, which counts the number of differences in the topology of the trees or the symmetric difference between trees. At 42.0, this value is high, suggesting many disagreements. In the chloroplast whole genome phylogeny, all but 3 nodes showed high support (LBS>65). This tree clusters *C. calisaya* specimens, except from Ccalisaya67 and Ccalisaya48, which cluster with *C. pubescens* and *C. nitida*, respectively. Similarly, we report the target capture supermatrix tree whose topology was mostly highly supported, with bootstrap values that were generally above 90. However, in this tree *C. calisaya* specimens cluster in other groups, for example with *C. pubescens*, that were not clustering in the plastid tree.

**Fig. 4.**
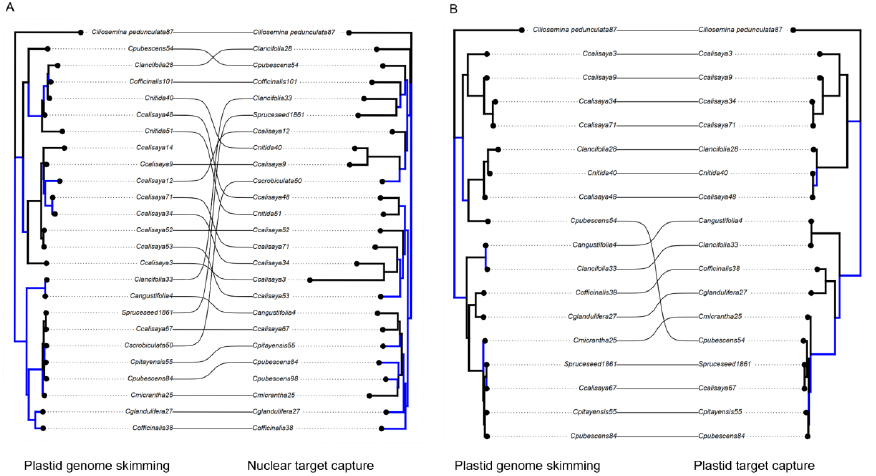
Tanglegrams showing, from left to right, (A) a comparison between the plastid genome skimming supermatrix and nuclear supermatrix tree based on target capture data. (B) The comparison of the plastid genome from genome skimming technologies with the plastid genome from target capture data. Blue highlights the disagreeing branches for both phylogenies, while black the branches that agree.

Since the nuclear and chloroplast genomes can have different evolutionary histories (e.g. due to hybridisation or incomplete lineage sorting) that likely underlie some of the discordance between the phylogenies produced by the two methods (genome skimming targeted the chloroplast and target capture nuclear loci), plastid reads were retrieved from the capture data. This was possible for 18 taxa, and a tree was estimated from these reads (Fig. 4B). Phylogenies were then compared without this confounding effect. The two trees using chloroplast data only were considerably more similar than the phylogenies targeting the different genomes (Fig. 4), with only a single taxon differing in placement (Cpubescens54).

## Discussion

Our principal aim was to demonstrate that robust and reproducible genomic characterisations can be produced from historical woody specimens, using the alkaloid-rich leaf and bark material from *Cinchona* species. By optimising the DNA extraction, library preparation, and bioinformatic pipelines, millions of reads were obtained from most of the specimens, succeeding in this objective (for 26 historical and six modern samples for each approach). We sampled 30 historical barks, of which 26 had high enough DNA quality and quantity to be sequenced. Three historical specimens failed to generate enough reads in each approach, two were the same samples that failed in both methods. This suggests that the main limitation to generating sequencing data for these samples was poor input quality DNA. There were also two different specimens that failed in each method, suggesting similar rates of success for each approach.

A secondary aim was to test the performance of the standard approach of genome skimming sequencing versus a new alternative, a standardised target capture kit (Angiosperms353) using these specimens. Here, the target capture approach performed generally better on the historical specimens, generating more raw reads and reads mapped (Fig. 2). There was however, no economic advantage to using the target capture approach since the savings from the superior sequencing depth were offset by the cost of purchasing the RNA baits required for library preparation. We therefore conclude that both methods produced enough genomic data to continue with downstream phylogenetic analyses in historical barks.

Plastid genome phylogenetics is the standard method for analysing herbarium material, because of the high copy number of the plastid genome. With the genome skimming approach, we were able to reconstruct the plastid genome using a reference-based approach, with a coverage up to 125x. However, this approach has the considerable limitation of failing to detect major genomic events that are only detectable within the nuclear genome (such as hybridisation and genome duplications). For example, hybridisation is common among the *Cinchona* species, especially within *Cinchona pubescens* (Andersson, 1998). *Additionally, the plastid genome has a limited impact in fully unravelling evolutionary processes in this group. For example, de novo* assembly is challenging because of the mostly ultra-short, damaged DNA molecules characteristic of *post-mortem* damage. Additionally, deeper sequencing does not necessarily result in higher coverage, because the endogenous DNA content can be very low in historical samples (Seitz and Nieselt, 2017). Thus, as reference databases are becoming more complete for non-model species, computational demands for analyses will decrease. Furthermore, as sequencing and bait synthesis costs continue to fall, full plastid genome reconstruction coupled with nuclear and plastid enriched loci could be the norm for future studies.

Target capture represents a powerful tool for analysing historical specimens that can capture major nuclear genome reconstruction events such as hybridisation and genome duplication. However, custom bait design is laborious and resource intensive. The recent standard ‘out of the box’ Angiosperms353 kit produced a larger number of raw and mapped reads. The previously outlined Paleomix pipeline (Schubert et al., 2014), retrieved a nominal number of reads for our target capture analysis. However, through the use of HybPiper, we recovered 108 genes on average for the historical samples and 160 for the silica dried modern samples. This suggests that with slight modification, the kit can be used for obtaining hDNA to infer phylogenies. Although previous studies have analysed herbarium samples with baits, including hybridisation processes (Larridon et al., 2019; Shee et al., 2020; Gardner et al., 2021; Brewer et al., 2019; Forrest et al., 2019), to our knowledge this is the first study that explores the ability of this technique to infer phylogenetic relationships using highly-degraded historical bark material. Using baits, many target loci within the nuclear genome can be simultaneously sequenced through the selective exclusion of non-target regions. However, it was possible to also retrieve high-copy regions, such as the chloroplast, from the target capture data. Additionally, multiple samples can be multiplexed within single reactions, although it is possible that multiplexing samples into single reactions reduces the ability of the baits to hybridise, and therefore reduce the quality of resulting data. Despite having the same number of samples in each lane, there was no evidence of ‘over-sequencing’ the samples in the target capture approach. Further increasing multiplexing of samples per sequencing run would likely negatively impact upon the number of loci that would be recovered. However, in a previous study of ragweed herbarium material, diluting custom baits to 10% of standard bait concentrations led to only a moderate reduction in read recovery (approximately ∼75%; Sánchez Barreiro et al., 2017; Hale et al., 2020), representing an alternative approach to decrease the additional costs of performing target capture.

Our final aim was to estimate the evolutionary relationship between samples using each method, and then compare them in terms of phylogenetic robustness and placement of taxa. Phylogenetic inference was performed using the genome skimming approach produced trees with very high branch support, and samples clustered more with their historical species annotation compared to the tree produced via target capture. When comparing the performance of both data types and approaches, nine tips out of 25 tips differed in locations (Fig. 4A). The discordance can nearly all be attributed to the different genomes being analysed, since only a single taxon differed in placement between the trees when chloroplast data was mined from both capture and genome skimming data (Fig 4B). Within the plastid trees, historical and modern *C. calisaya* specimens clustered together more than within the nuclear tree, with the exception of the samples Ccalisaya48 and Ccalisaya67. While the nuclear supermatrix analysis from the target capture approach shows that modern *C. calisaya* specimens (Ccalisaya52 and Ccalisaya53) cluster with two historical ones (Ccalisaya71 and Ccalisaya34), and the remainder of historical *C. calisaya* specimens are distributed across the tree within different clusters, e.g. with a *C. pubescens* cluster that includes Cpubescens84 and Spruceseed1861. Discordances were likely a consequence of either inaccurate historical annotations or large-scale genome reorganisation events. Using well-annotated samples or type specimens can give an accurate perspective on the analysed genetic data.

The ability to produce detailed genomic characterisations from historical collections would allow for us to determine unknown or contested species designation, unravel historical trade routes, assess conservation status, and shed light on the origin of bred samples. Even within *Cinchona*, a genus of considerable importance within the pharmaceutical field and the beverage industry, there is still much to be learnt about the genus’ biological and historical processes. For example, there is currently no reference nuclear genome (or transcriptome) within the tribe, making the design of custom baits challenging, and emphasises the value of ‘out-of-the-box’ kits such as the Angiosperms353 kit. However, biocultural collections pose a number of challenges. Many historic collections are annotated incompletely, missing metadata such as species, dates, and origins. Within the case of *Cinchona*, specimens are sometimes listed under trade names rather than species, and labelled under the cities they were exported from, rather than the true sampling location. In addition, they may also have been mislabelled historically. This may have been unintentional and deliberate adulteration, since there were economic incentives to mislabel the barks of other less valuable species as *Cinchona*. This poses a challenge since the annotated specimens cannot be reliably used as a reference for the phylogenetic studies. The discordance between our trees and the potentially adulterated bark specimens make it uncertain which method showed more accurate evolutionary relationships. Modern phylogenetic studies in Rubiaceae have been able to assign near to species level, thus big differences between the topologies are not likely to happen (Manns and Bremer, 2010). Within the samples used within this study (RBGK), it would be possible to determine the provenance of samples using both approaches performed here, and similarly, they could be implemented on other highly degraded herbarium material like wooden artefacts, and samples of significant biocultural heritage.

## Conclusion

Our study demonstrates the utility of highly degraded plant material, such as historical bark specimens, to conduct large-scale palaeogenomic studies using target capture. Based on our results, we recommend the target capture Angiosperms353 kit as a powerful and robust method inferring nuclear and plastid phylogenies from highly degraded samples. However, this is for taxa without a reference genome where reads cannot be mapped. If genomic resources are available (transcriptome), it should be considered whether the time and resources should be allocated to produce custom baits that would almost certainly yield optimal results.

## Acknowledgements

The authors thank Stefanie Wagner, Christian Carøe, Fátima Sánchez-Barreiro, and Thibauld Michel for their input with the DNA extraction and library building protocols. To Rafal Gutaker for his help discussing this study. To Lasse Vinner for sequencing strategy support. The authors thank the GeoGenetics Sequencing Core for assistance in generating the Illumina data.

## Data Accessibility

All sequencing data for both genome skimming and target capture will be made freely available on the Dryad repository upon manuscript acceptance.

## Author Contributions

CB and NC conceived and designed the study. NC performed the sampling with input from KW and MN. NC performed the laboratory work with help from TG. NC and CB performed the statistical analyses. NC performed the computational analysis with input from EMG, VB, MDM, and CB. KW and MN performed the archival search. NC wrote the manuscript with input from CB and all authors.

## Supporting material

Appendix Fig. S1. *Post-mortem* damage of a historical bark *Cinchona* sample (Ccalisaya14).

Appendix Fig. S2 Plastid genome skimming phylogenetic tree and bootstrap values of historical and modern *Cinchona* samples.

Appendix Fig. S3 Nuclear supermatrix target capture phylogenetic tree and bootstrap values of historical and modern *Cinchona* samples.

Appendix Tab. S1. Overview of specimens and sample preparation for this study.

Appendix Tab. 2. Advantages and disadvantages of using genome skimming and target capture on historical barks.

## Data accessibility statement

Historical and modern DNA genomic data sets, gene recovery and mapping files will be deposited in Dryad upon acceptance of the manuscript.

