## Supplementary figures and images for "Museomic approaches to genotype historic *Cinchona* barks"

### Supp. fig 1

#### mapDamage plot for library 'Ccalisaya14'

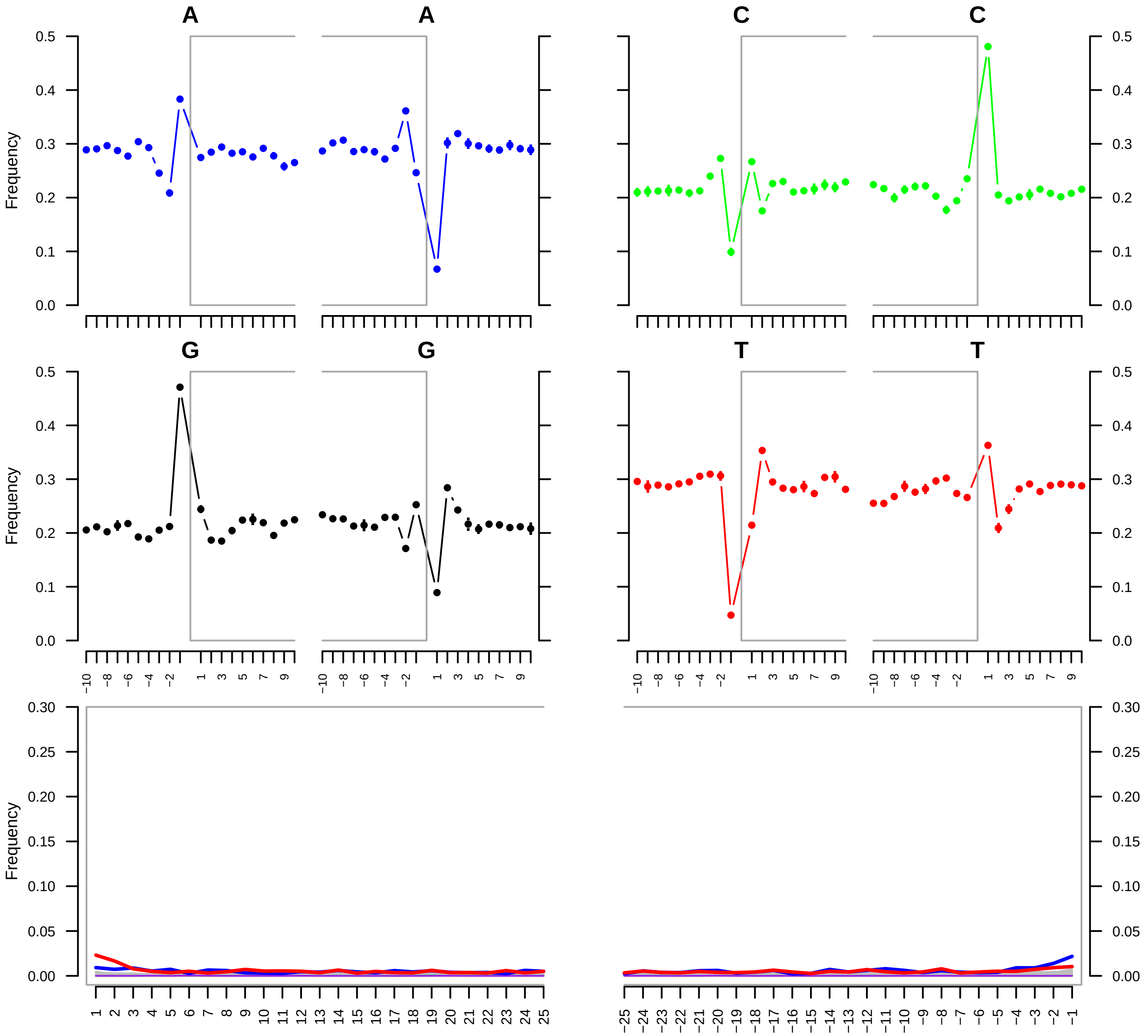

### Supp. fig 2

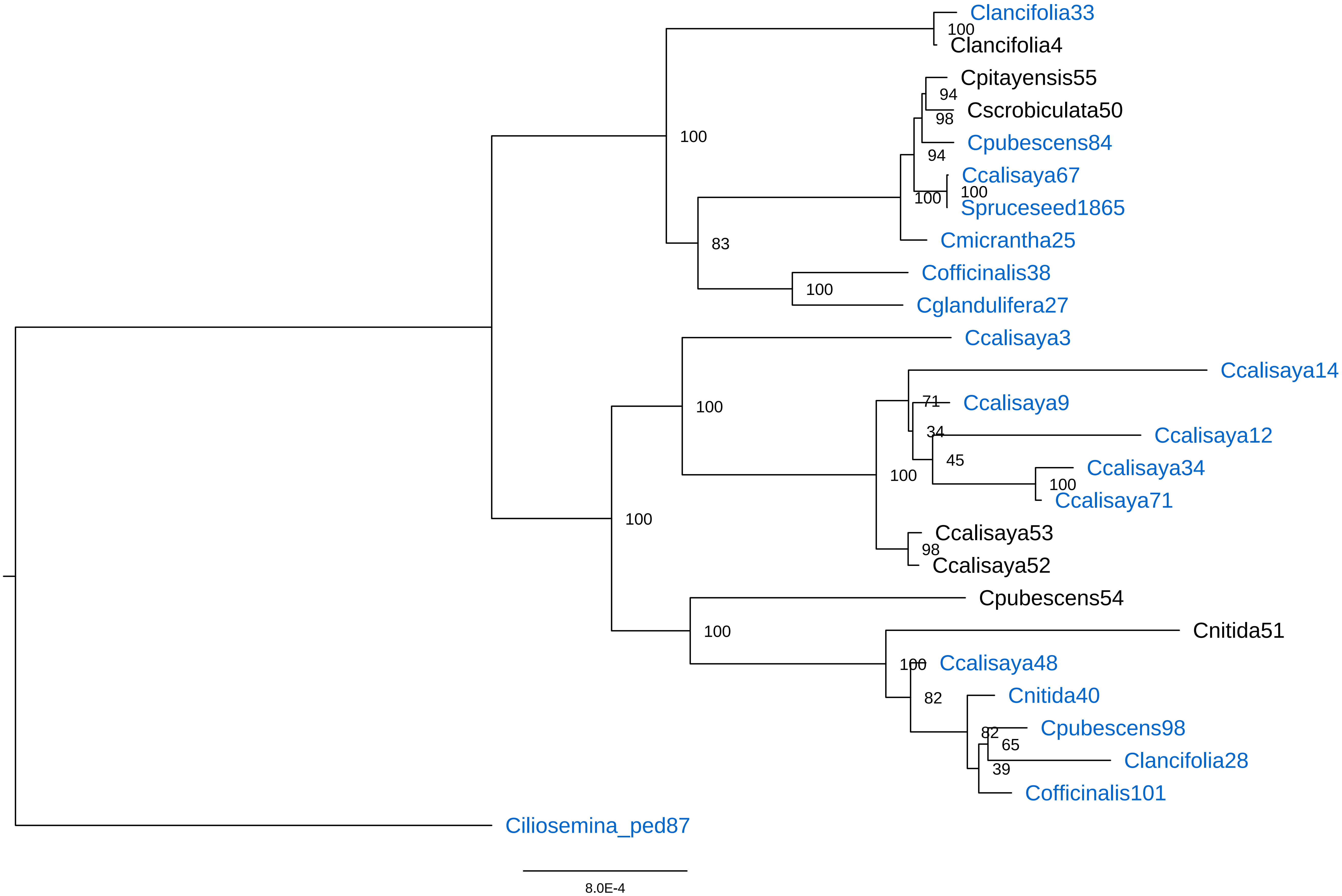

### Supp. fig 3

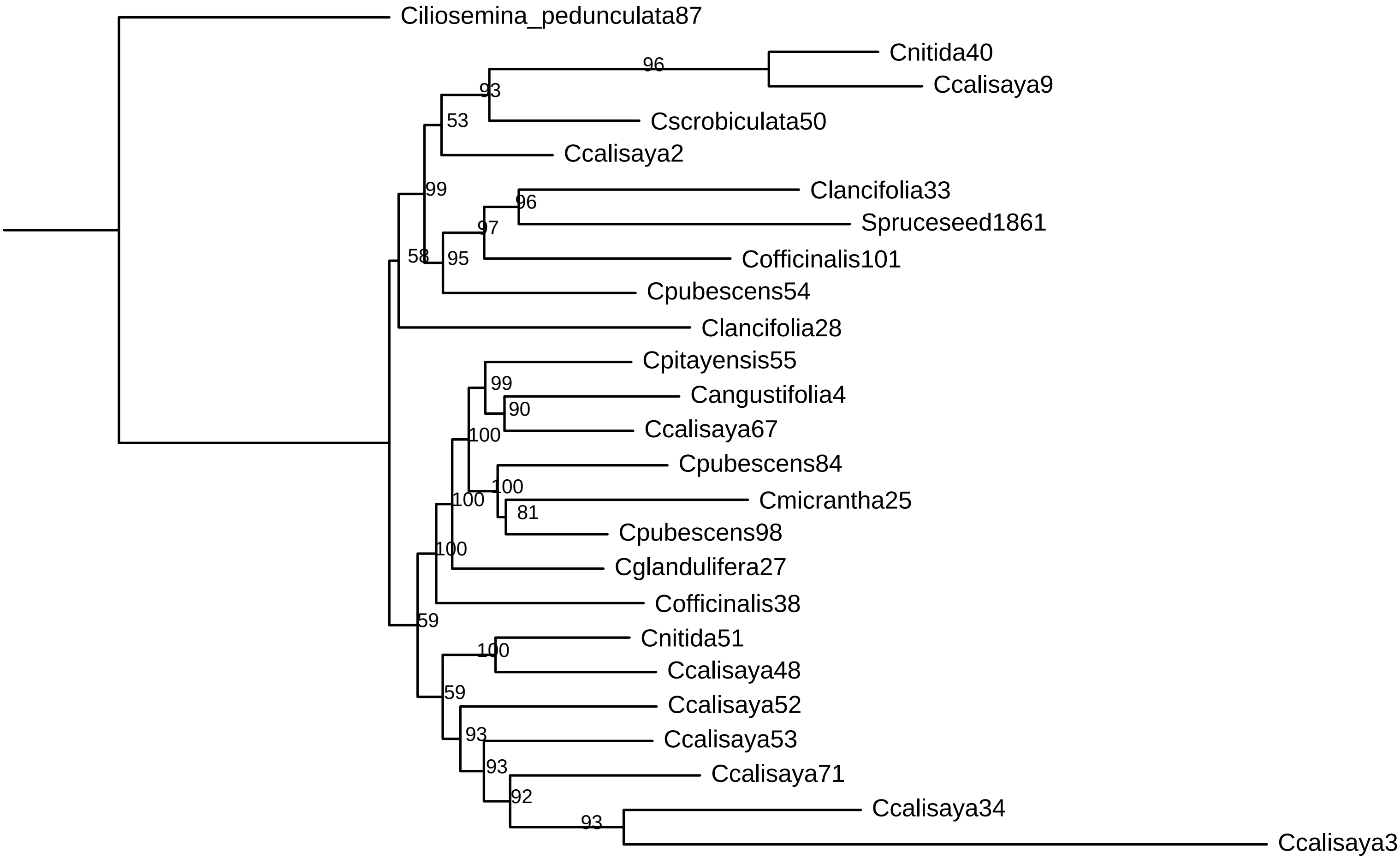
